# Latency structure of BOLD signals within white matter in resting-state fMRI

**DOI:** 10.1101/2021.03.06.434206

**Authors:** Bin Guo, Fugen Zhou, Muwei Li, John C. Gore

**Affiliations:** Image Processing Center, School of Astronautics, Beihang University, Beijing 100191, Beijing, China; Vanderbilt University Institute of Imaging Science, Vanderbilt University, Nashville, TN 37232, USA; Department of Radiology and Radiological Sciences, Vanderbilt University Medical Center, Nashville, TN 37232, United States; Department of Biomedical Engineering, Vanderbilt University, Nashville, TN 37232, United States; Department of Molecular Physiology and Biophysics, Vanderbilt University, Nashville, TN 37232, United States; Department of Physics and Astronomy, Vanderbilt University, Nashville, TN 37232, United States

**Keywords:** BOLD, fMRI, functional connectivity, resting state, white matter

## Abstract

**Purpose:** Previous studies have demonstrated that BOLD signals in gray matter in resting-state functional MRI (RSfMRI) have variable time lags, representing apparent propagations of fMRI BOLD signals in gray matter. We complemented existing findings and explored the corresponding variations of signal latencies in white matter.

**Methods:** We used data from the Brain Genomics Superstruct Project, consisting of 1412 subjects (both sexes included) and divided the dataset into ten equal groups to study both the patterns and reproducibility of latency estimates within white matter. We constructed latency matrices by computing cross-correlations between voxel pairs. We also applied a clustering analysis to identify functional networks within white matter, based on which latency analysis was also performed to investigate lead/lag relationship at network level. A dataset consisting of various sensory states (eyes closed, eyes open and eyes open with fixation) was also included to examine the relationship between latency structure and different states.

**Results:** Projections of voxel latencies from the latency matrices were highly correlated (average Pearson correlation coefficient = 0.89) across the subgroups, confirming the reproducibility and structure of signal lags in white matter. Analysis of latencies within and between networks revealed a similar pattern of inter- and intra-network communication to that reported for gray matter. Moreover, a dominant direction, from inferior to superior regions, of BOLD signal propagation was revealed by higher resolution clustering. The variations of lag structure within white matter are associated with different sensory states.

**Conclusions:** These findings provide additional insight into the character and roles of white matter BOLD signals in brain functions.

## 1. Introduction

Functional magnetic resonance imaging (fMRI) studies were initially based on the detection of changes in blood oxygenation-level dependent (BOLD) signals after a task or stimulus. The technique was later extended to include the detection and analysis of spontaneous signal fluctuations [1]. Though initially regarded as noise, these spontaneous fluctuations were later analyzed to reveal intrinsic synchronizations of regions within the somatomotor system of the brain [2], and these result in long range, inter-regional low frequency temporal correlations in measured BOLD signals. Conventional analyses of these temporal correlations yield a metric widely referred to as functional connectivity (FC) which is derived by assuming exact synchronization of activities and BOLD signals in intrinsic brain networks. However, previous studies in rat and humans found that BOLD fluctuations are spatio-temporally organized across different brain circuits [3][4]. To characterize the spatiotemporal patterns of brain activities, Mitra and colleagues recently analyzed inter-regional time latencies or lags in BOLD signals. They derived a time delay matrix from those lags corresponding to extrema in the cross-correlation functions of pairs of voxels. A highly reproducible, state-dependent latency projection (the summed lags relative to other voxels) spanning around 1 second, was found to correspond to spatially segregated functional networks [5][6]. Since then, much effort has been paid to comprehensively explore lead/lag relationships within the cerebral cortex [5], between the thalamus and the cerebral cortex [7], and the cerebellum [8]. Such relationships are important for appropriate data analysis and for the interpretation of functional interactions within and between networks.

In contrast to studies of BOLD signals in gray matter, BOLD signals in white matter have been ignored for most of the past 30 years and their existence has even been regarded as controversial. In many analyses of fMRI data from gray matter, signals within white matter are treated as nuisance regressors. However, recent growing evidence converges to support the conclusion that fluctuations in white matter BOLD signals encode neural activities [9][10][11], similar to gray matter. It is not clear whether BOLD signals in white matter reflect an intrinsic metabolic demand or are driven by activity in grey matter. Several studies have illustrated the characterisics of BOLD signals in white matter. For example, BOLD signals are robustly detectable in WM if appropriate analyses are used; conventional GLM methods throw away sensitivity by using an inappropriate hemodynamic response function (HRF) [11][12]. Event related measurements show the HRF is different from GM but can be measured accurately [13][14]. WM BOLD activity can be evoked by stimulation in task-specific tracts or regions. The magnitude of WM signals in a task reflects those in GM engaged in the same task, and may be modulated by the same factors [15][16]. At rest, WM tracts show reproducible patterns of connectivity which are summarized in Functional Connectivity Matrices (FCMs) obtained by analyzing resting state correlations between segmented WM and GM parcellations [17]; the FCM relating WM to GM is altered in various pathologies including Alzheimer’s disease in a manner that correlates with behavioral measures [18]. Distinct, reproducible networks in WM emerge from data-driven analyses in similar manner to cortical circuits [19]. Moreover, comparisons of partial and full correlations between GM regions with inclusion vs exclusion of WM shows the degree of engagement of specific WM tracts in the couplings between cortical volumes [20]. Peer and colleagues investigated the functional networks of white matter by applying simple clustering analysis on RSfMRI data of white matter regions [21]. Highly bilaterally symmetrical functional networks were identified. These and other studies suggest white matter BOLD signals should be incorporated into models of functional networks.

Despite this converging evidence for the functional role of BOLD signals in white matter, temporal properties of these signals have not been adequately investigated. White matter connects various gray matter areas of the brain to each other, and is responsible for transducing neural activity between regions. In light of recent advances in modelling of how networks communicate [22], examination of the timing relationships within white matter is particularly compelling.

Here we report studies of the latency structure of BOLD signals within white matter. The results are derived from the Brain Genomics Superstruct Project (GSP), a publicly available dataset consisting of a large cohort of healthy young subjects. We found that a highly reproducible latency projection also exists in white matter. Further latency analysis of inter- and intra-functional networks, identified by clustering on functional connectivity, revealed that some of the functional networks within white matter also exhibit similar spatiotemporal organizations to those in gray matter [5]. Moreover, using the BeijingEOEC dataset II, we found that variations of the lead/lag relationships within white matter were associated with different sensory states of the brain, suggesting a causal effect between them.

## 2. Methods

### 2.1 Subjects and MRI acquisitions

Resting-state data from 1412 healthy subjects (both genders included) were selected from the Brain Genomics Superstruct Project (GSP) [23]. All imaging data were acquired on matched 3T Tim Trio scanners (Siemens Healthcare, Erlangen, Germany) at Harvard University and Massachusetts General Hospital using vendor-supplied 12-channel array head coils. The data for each subject consisted of a 6-minute resting-state echo-planar imaging (EPI) scan with eyes open (38 slices, TR=3s, 3×3×3 mm isovoxels, interleaved slices) and an MPRAGE anatomical scan (1.2×1.2×1.2 mm resolution). Participants involved provided written informed consent in accordance with guidelines established by the Partners Health Care Institutional Review Board and the Harvard University Committee on the Use of Human Subjects in Research [23].

We further used resting-state data from 20 healthy subjects (11 females) in the BeijingEOEC dataset II [24][25] to investigate the influence of different sensory states on the lag structure in brain white matter. The following three resting-state sessions were acquired and counterbalanced across the participants: 1) EC (eye closed), 2) EO (eye open), and 3) EOF (eyes open with a fixation). Each of the sessions lasted for eight minutes. During the three resting-state sessions, the participants were instructed to keep as motionless as possible and to think of nothing except for mind wandering. During the EOF session, the participants were instructed to fixate on a black crosshair in the center of a white screen. The functional images were obtained using an EPI sequence with the following parameters: 33 axial slices, thickness=3 mm, gap=0.6 mm, in-plane resolution=64×64, TR=2000 ms, TE=30 ms, flip angle=90°, FOV=200×200 mm. In addition, a T1-weighted sagittal three-dimensional magnetization-prepared rapid gradient echo (MPRAGE) sequence was acquired, covering the entire brain: 128 slices, TR=2530ms, TE=3.39ms, slice thickness=1.33 mm, flip angle=7°, inversion time=1100 ms, FOV=256×256 mm, and in-plane resolution=256×192. This study was approved by the ethics committee of state key laboratory of cognitive neuroscience and learning, Beijing Normal University. Written informed consent was obtained from each subject.

### 2.2 Functional and anatomical preprocessing

All images were preprocessed using the statistical parametric mapping software package SPM12 (www.fil.ion.ucl.ac.uk/spm/software). The initial four time points of each BOLD series were discarded to allow for equilibrium to be reached. First, the images were corrected for slice timing and head motion, and subjects with large head motion (maximum translation >2 mm or maximum rotation >2°) were excluded, resulting in a cohort of 1412 subjects. Second, T1-weighted images were segmented into gray matter, white matter, and cerebrospinal fluid using the New Segment utility embedded in SPM12, and all these images were registered to the mean BOLD image output by the motion correction procedure. Third, the BOLD images were normalized into the Montreal Neurological Institute (MNI) space, along with the coregistered T1-weighted images as well as the gray matter and white matter segments. Fourth, linear trends (first order) from the BOLD images were removed to correct for signal drifts using the detrend function in matlab. Fifth, mean signals from the cerebrospinal fluid mask and whole-brain mask, and also Friston’s 24 regressors [26] were regressed out as nuisance covariates. After the nuisance regression, the temporal signals were low-pass filtered (using FFT) to retain frequencies between 0.01∼0.1Hz. Finally, BOLD images were spatially smoothed with an isotropic Gaussian filter (FWHM = 6 mm), where smoothing was performed separately on the white matter and the gray matter of each subject, to avoid mixing of signals.

### 2.3 Creation of group-level white matter mask

Following the preprocessing, the white matter mask of each subject was normalized to the MNI space and resampled to isotropic resolution of 3 mm. To obtain the masks for group-wise analysis, we used the segments for each subject. Each subject-level white matter mask was generated using a threshold value of 0.95, and the strategy of majority voting was applied to generate the group-level mask. Voxels with a voting rate above 90% were included resulting in a group-level mask consisting of 9321 white matter voxels.

### 2.4 Latency analysis

We followed the method proposed by Mitra et al. [32] for our latency analyses. A schematic diagram of our latency analysis for this study is shown in Figure 1. Details are included in the following sub-sections.

**Figure 1.**
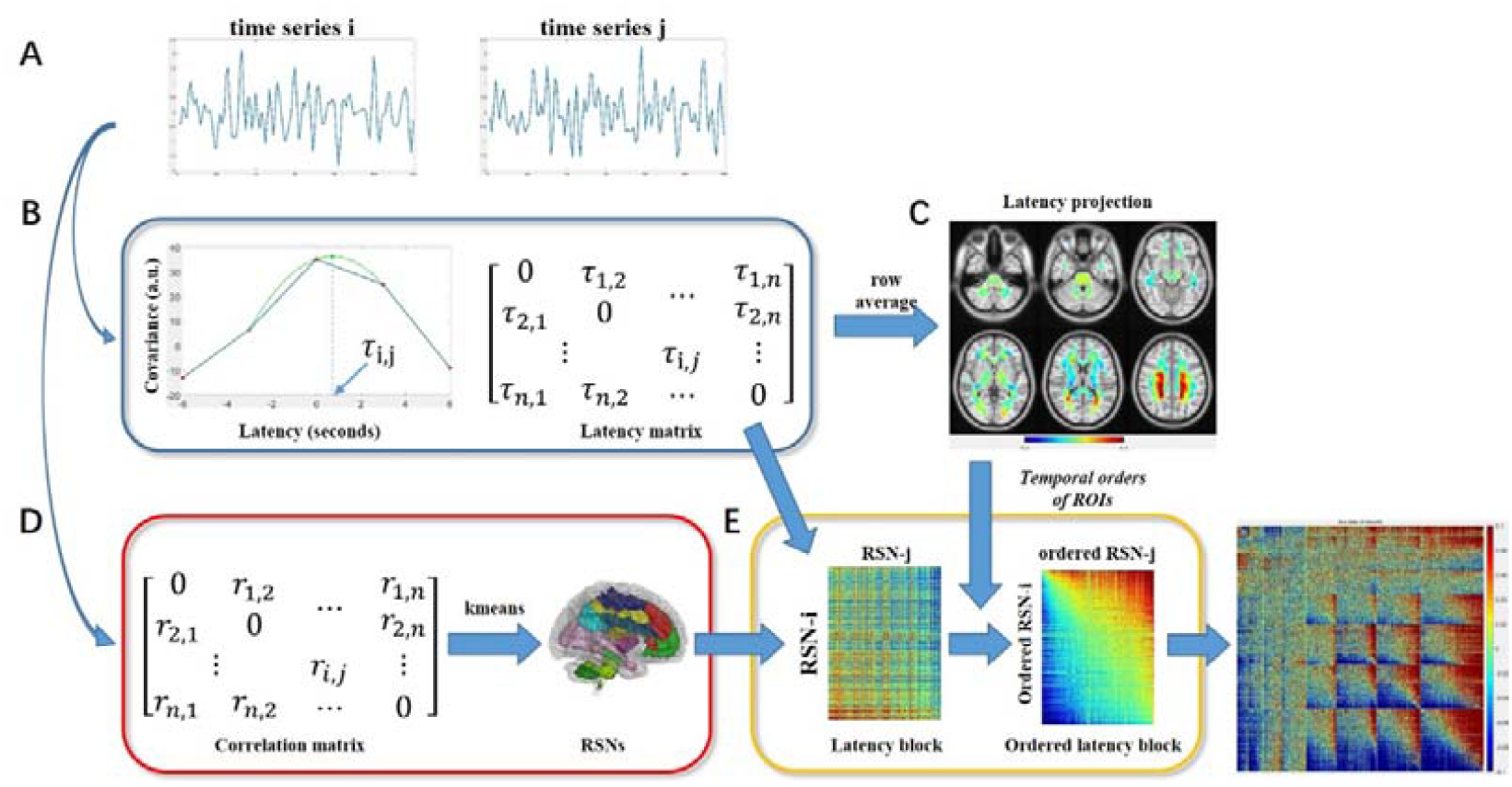
Schematic illustrating the workflow of latency analysis. (A) Time series of two ROIs, i and j. An ROI could be a single voxel or a submask of a WM mask. Mean time series will be used if a submask was selected as an ROI. (B) Estimation of lag value of two ROIs. Lagged cross covariance was computed first, followed by parabolic interpolation to determine the latency value corresponding to maximum cross covariance. Latencies from all paired ROIs constitute the latency matrix, which is necessarily anti-symmetric. (C) Latency projection was obtained by averaging along the rows of the latency matrix. The value of each ROI was mapped back to the original WM mask. Temporal orders or each ROI can be derived from the latency projection. (D) Computation of correlations among all ROIs results in a symmetric correlation matrix. A Kmeans clustering procedure was applied on the group-level correlation matrix to obtain RSNs in WM. (E) The RSNs were utilized to group entries in the latency matrix into latency blocks. These latency blocks were ordered by incorporating the temporal orders derived from the latency projection, wherein early ROIs were placed near the left and top corner while late ROIs the right and bottom corner of the latency block. Repeating this operation yielded an ordered latency matrix at the RSN-level, in which off-diagonal blocks represented inter-RSN lead/lag relationship and diagonal blocks the intra-RSN lead/lag relationship.

#### 2.4.1 Generation of subject- and group-level correlation matrix

The time series of each voxel in WM mask were extracted. For each subject, Pearson’s correlation was computed for all pairs of voxels, resulting in a subject-level correlation matrix of size 9321×9321. A group-level correlation matrix was defined to be the average of all the subject-level correlation matrices within the same group.

#### 2.4.2 Construction of white matter functional networks

Functional networks of white matter were identified by clustering voxels on the basis of group-level correlation matrices generated as above [27][28][29][30][31]. K-means clustering with correlation as the distance metric and with twelve replicates was performed on the rows of each group-level correlation matrix, which allowed voxels with similar functional connectivity patterns to be grouped into the same cluster. In this study, K-means clustering was carried out for each subgroup, as well as for the entire dataset by pooling all the subjects together, which served as ground truth.

The cluster similarity between each subgroup and the ground truth was evaluated. We first computed an adjacency matrix, wherein each element in the matrix indicates whether any two voxels are in the same cluster or not, for each clustering result [31]. We then computed the Dice coefficients between the adjacency matrices from each subgroup and from the entire dataset.

#### 2.4.3 Estimation of time delay between two time series

Briefly, given time series of two voxels, *X*_1_(t) and *X*_2_(t), a lagged cross-covariance was computed as,

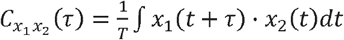

where τ is the lag (in units of time), and T is the integration interval, a constant in our experiment. The value of τ at which 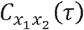exhibiting an extremum defines the temporal lag (delay) between signals *X*_1_ and *X*_2_. To determine the lag time value at a temporal resolution finer than the time of repetition, a parabolic interpolation was used, as described previously [5][34].

#### 2.4.4 Generation of time delay matrix and latency projection

A latency matrix was computed for each subject and then averaged across each group to obtain group-level latencies, a matrix of size 9321×9321. A latency projection was then obtained by averaging over the rows of the latency matrix. The latency projection represents the relative timing of each voxel with respect to the remaining voxels within white matter.

#### 2.4.5 Generation of latency matrices among networks

To characterize the lead/lag relationships among networks, rows and columns of the latency matrix were first rearranged by clusters such that voxels of the same cluster were adjacent to each other, thus subdividing the latency matrix into K×K blocks. Second, intra-network voxels were reordered by their temporal orders derived from latency projections. Thus, diagonal blocks represent the lead/lag relations between intra-cluster voxels and off-diagonal blocks the lead/lag relations between inter-cluster voxels. Within each block of cluster pairs, the voxels with early timings relative to the remaining voxels in the same block were placed in the upper-left corner and voxels with later timings were placed in the lower-right corner.

#### 2.4.6 State contrast of the latency structure within white matter

We examined statistical differences in the lag structure within white matter among the following three different resting sensory states, i.e., EO, EC, and EOF. A linear mixed-effects analysis [35] was adopted to study the main effect of the sensory states, along with the in-between paired effects. The modeling program, 3dLMEr from AFNI [36], was used to build the statistical model, in which variates of sensory states (EO, EC, and EOF), age, and gender were included. Cluster correction was employed for evaluating both main and paired effects. Clusters of statistical significance were identified based on their size. The threshold of cluster size regarded as statistically significant was determined by autocorrelation function modeling combined with a voxel-level threshold of p<0.001. Specifically, we applied 3dFWHMx on preprocessed fMRI data to estimate the parameters of the autocorrelation function (ACF), followed by 3dClustSim for identification of significant cluster-size, within which a corrected significance level was thresholded at p<0.05 and uncorrected voxel-level at p<0.001 [37].

## 3. Results

### 3.1. Latency projections in white matter

Latency projections within WM derived from the GSP dataset are shown in Figure 2 wherein positive values indicate lags and negative values leads. This lead/lag relationship appears as signal propagations from the leads to the lags, thus *apparent propagation*. It can be seen that the distribution of the latencies was highly bilaterally symmetric. The white matter regions with the smallest and greatest lag values were the internal capsule (IC) and the posterior corona radiate (PCR), respectively. A gradual increase in lag value from the posterior thalamic radiation (PTR) to the optic radiation (OR) was also observed. The correlogram in Figure 2-B shows the pair-wise correlation coefficient among the ten separate subgroups. The correlation value ranged from 0.859 to 0.908, which demonstrates high spatial similarities in latency projections across subgroups.

**Figure 2.**
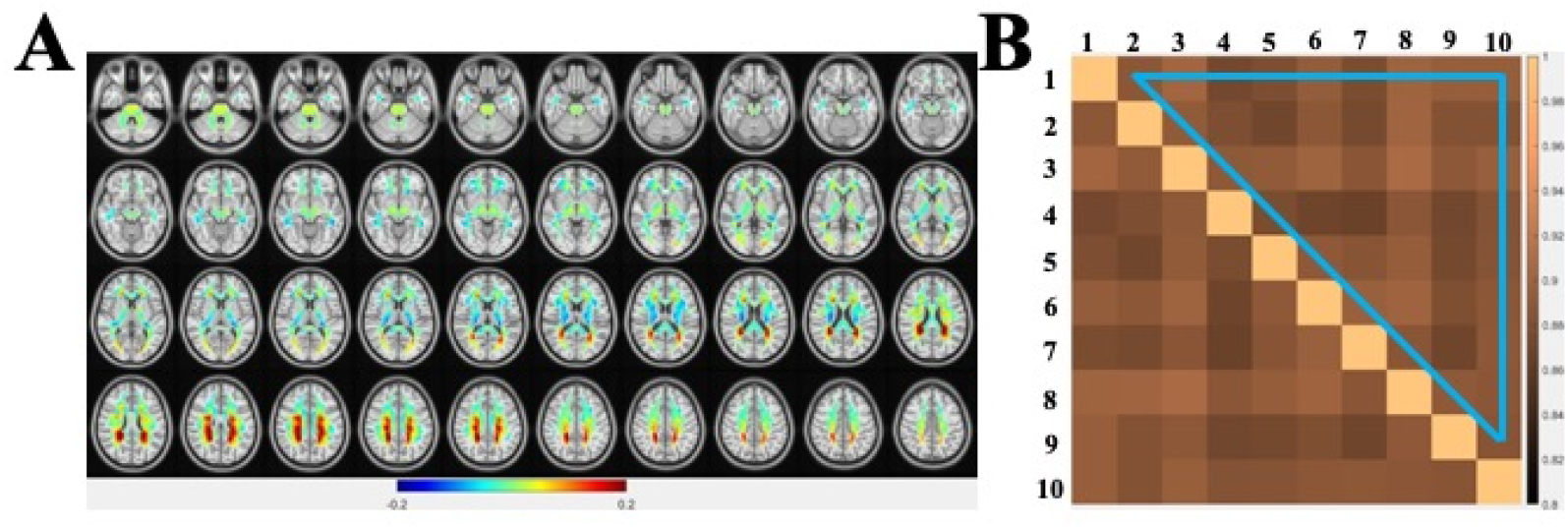
High reproducible latency projection from one group of the ten equal size groups. (A) Voxel-wise latency projections of TD obtained from the first group within white matter. (B) Pair-wise spatial correlations in latency projections between the ten subgroups studied. The mean and standard deviation within the upper triangular region, demarcated by the blue triangle, of the correlogram are 0.887 and 0.013, respectively).

### 3.2 Functional networks and latency matrix

#### 3.2.1 Functional networks within white matter

Functional networks within white matter identified by using Kmeans clustering are shown in Figure S1. The numbers of clusters that yield the most stable cluster segregations are 2, 4, 8, and 11 (denoted as asterisks in Figure S1-A). These clusters are visualized in left, right, front and back views in Figure S1-B.

#### 3.2.2 Latency matrices among white matter functional networks

We examined lead/lag relationships among white matter functional networks (WMFN) on different numbers of clusters in a coarse-to-fine manner. Here the latency matrices of only 2, 4, and 8 clusters are shown in Figures 3-4, with diagonal blocks representing intra-network lead/lag relationships, while off-diagonal blocks correspond to inter-network lead/lag relationships. Lead/lag relationships at the coarsest level (K=2) of clustering are shown in the left of Figure 3, in which the two networks clustered are in the top (green colored) and bottom (red colored) portions of the brain respectively. Note that a portion of the anterior corpus callosum was clustered into the bottom cluster. For the top cluster, a pattern ordering the intra-network latencies was readily identified (see yellow rectangle in Figure 3-A). This latency pattern resembles previous studies on functional networks of gray matter [5], indicating similar spatiotemporal properties such that top cluster appeared to contain temporally well-ordered early, medium and late components within WMFN. Meanwhile, for the bottom cluster, a reverse pattern to the top cluster’s intra-network lead/lag relationship was observed (see red rectangle in Figure 3-A), indicating a reverse temporal ordering derived from the latency projection against that derived from within cluster. For inter-network lead/lag relationship, the latency patterns in the off-diagonal blocks were quite different from those found in gray matter previously. Part of the early voxels from the top cluster lag part of the early voxels from the bottom cluster. Clusters with K=4 are shown in the right of Figure 3. The top cluster obtained when K=2 was further subdivided into two clusters, one covering the middle part of the original top cluster (named as TM, short for top-middle) and the other both ends of the original top cluster (named as TE, short for top-ends). Similarly, the bottom cluster obtained when K=2 was further clustered into top (BT, short for bottom-top) and bottom (BB, bottom-bottom) components. It can be seen that there were well-ordered distributions of inter- and intra-network latencies between network TM (blue) and network TE (yellow), indicating the existence of reciprocal inter- and intra-network timings (see yellow rectangle in Figure 3-D). At this clustering resolution, network BT (green) exhibited a clear reverse intra-network temporal order against the temporal order deduced from the latency projections (see red rectangle in Figure 3-D). In network BB (red), the overall temporal order was unclear though some of the voxels led some of the voxels from other clusters.

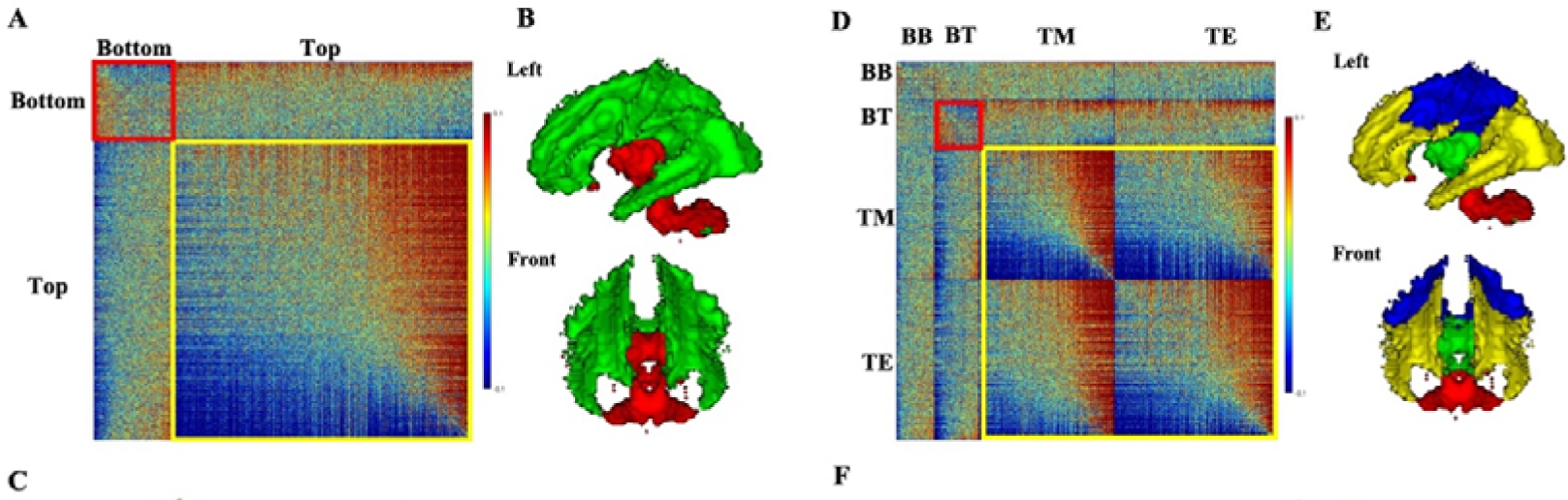

To establish anatomical correspondences, clusters were named according to the largest WM fiber bundle they contained, including middle cerebellar peduncle (MCP), posterior limb of inter capsule (PLIC), retrolenticular part of internal capsule (RPIC), corpus callosum (CC), anterior corona radiata (ACR), posterior thalamic radiation (PTR), superior longitudinal fasciculus (SLF), and superior corona radiata (SCR), which are shown in Figure 4-B. Latency analyses among these RSNs are displayed in Figure 4-A, wherein reciprocal inter- and intra-network propagations were observed among the lower-right blocks (see yellow rectangle in Figure 4-A), consisting of SCR, ACR, PTR, and SLF, but no temporal relationships were clearly visible between network MCP and other networks except RPIC. CC consisted of a major portion of the corpus callosum, the temporal relationships of which with MCP seemed disordered.

**Figure 4.**
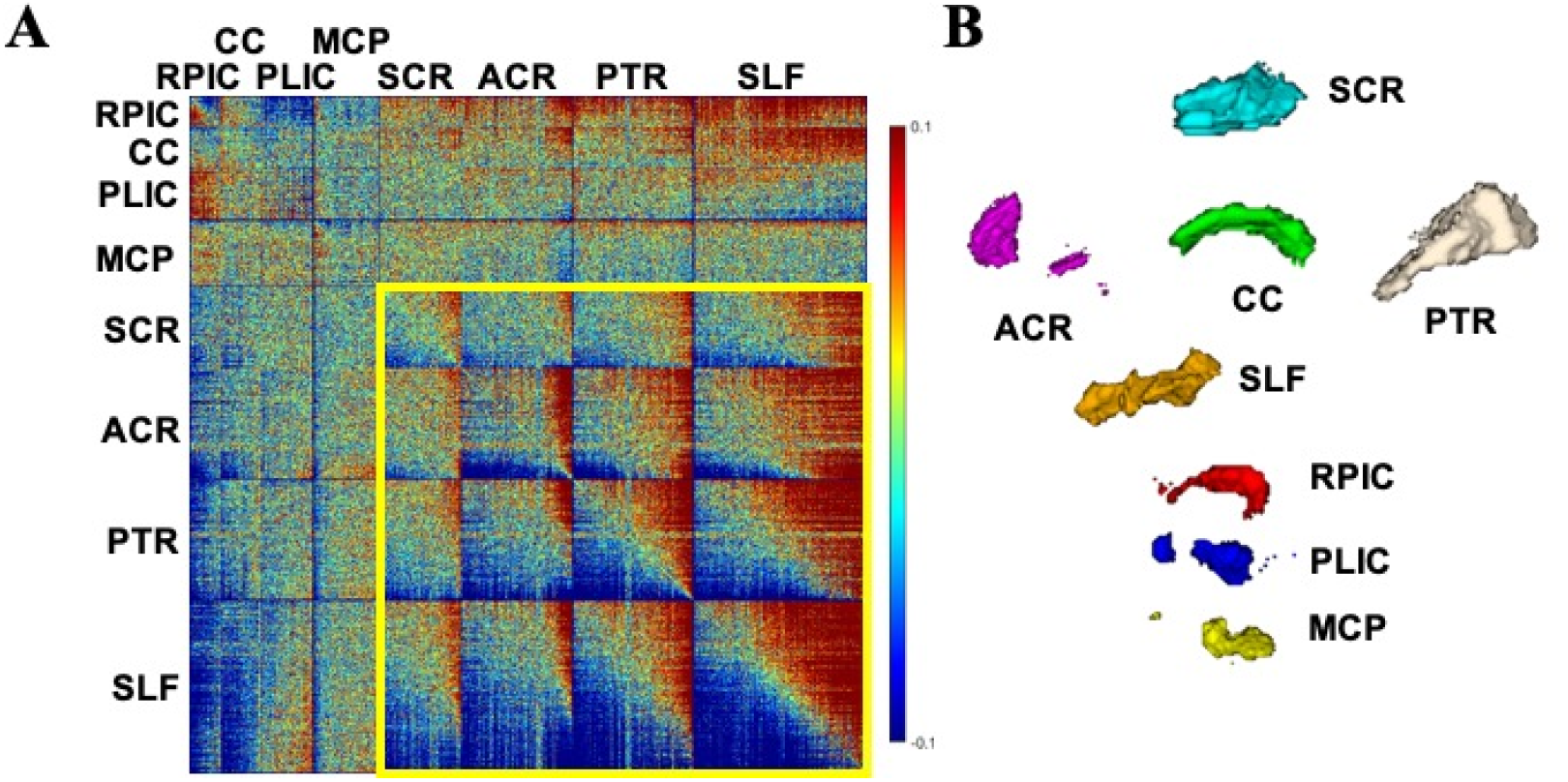
Latency matrix and clustering result when K=8. (A) Latency matrix. (B) Names of clusters. Clusters are arranged according to their spatial positions.

Contrasting with the results from GM [22] wherein no significant lead/lag relationships at the network level were identified between RSNs, latency analysis in WM revealed the existence of inter-RSN lead/lag relationship, as indicated by asterisks in Figure 5-A (one sample t-test on the mean value of each off-diagonal block, p<0.05 with Bonferroni correction, N=28), which suggested that propagations of BOLD signals between paired RSNs, indicated by each asterisk, appeared to be dominated by one direction. Thus, a dominant direction of BOLD signal propagation (indicated by black arrows while the other direction was indicated by blue arrows in Figure 5-B), from inferior WM (including PLIC) to middle WM (including RPIC) and further to superior WM (including PTR, SCR, SLF, and PTR) was identified. Also identified were the signal propagations among the RSNs in superior WM, including CC, PTR, SCR, SLF, and PTR, but with weaker directional dominance, as shown in the right of Figure 5-B. Latency matrices of other clusters are provided in Figures S2∼S7 in the Supplementary Material.

**Figure 5.**
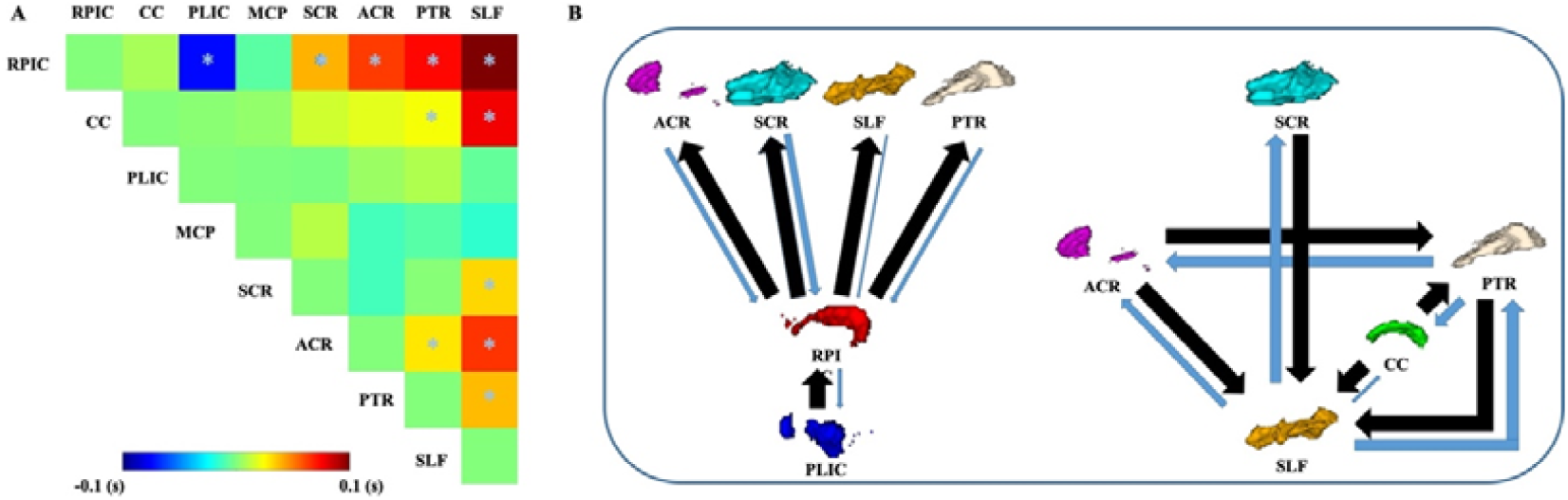
Identification of dominant paths of signal propagation. (A) Results of one-sample ttest. Blocks indicated by asterisks showed significant (p<0.05, with Bonferroni correction of N=28) non-zero mean lags, suggesting dominant direction of signal propagations. (B) Paths showing directional dominance. Black arrows indicate the dominant direction, i.e. most voxels from target (pointed) RSN lag those from source (originated) RSN, and blue arrows the other direction. Sizes of black arrows are fixed and sizes of the blue arrows are proportionated accordingly by the ratio of count of negatives over count of positives in each latency block.

### 3.2 Latency projections of different sensory states

Latency projections obtained during different sensory states are shown in Figure 6. As can be seen, the spatial patterns of latency projections in white matter were largely preserved across different sensory states, which were also reminiscent of those obtained from the GSP dataset in Figure 2. Pearson’s correlation analysis showed that the CC between EO and EC, EO and EOF, and EC and EOF were 0.798, 0.805 and 0.809 respectively. Close inspections revealed that, for all the three states, voxels with large lag values were mostly distributed in the corona radiata (CR), OR, and the four corners of ventricles, whereas voxels with small lag values were mainly concentrated around the IC. Notable differences in lag values among the three sensory states, however, were also clearly visible, especially in the IC and OR. Larger lag values tended to be concentrated around the OR for EC as compared with EO and EOF, while small lag magnitudes tended to be around the IC for EO and EOF.

**Figure 6.**
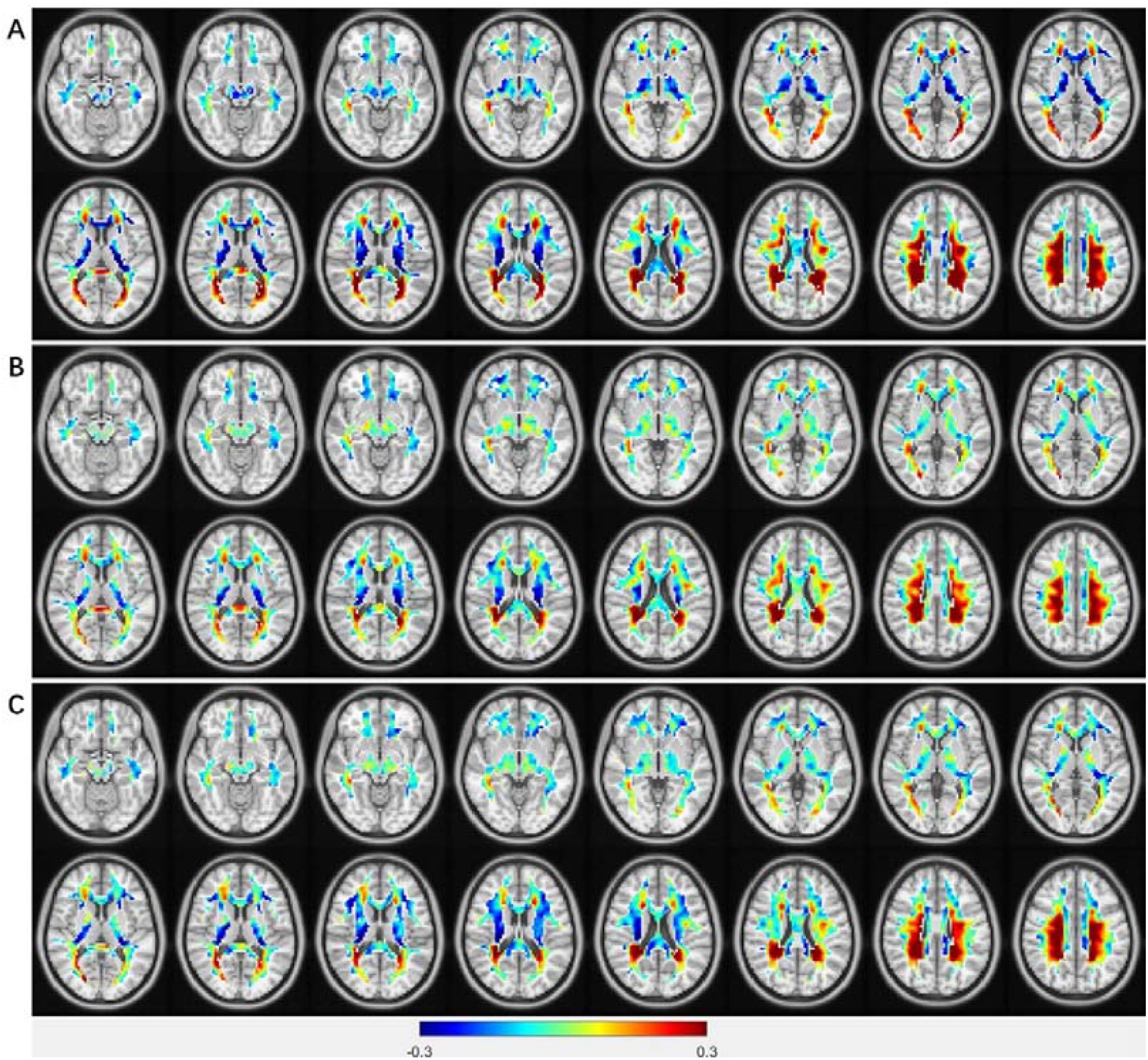
Latency projections in white matter obtained in the Beijing EOEC dataset II. (A) EC. (B) EO. (C) EOF. Note that the distributions patterns across the three different states are grossly consistent.

Figure 7 shows the regions exhibiting significant differences (defined by FWE-corrected p<0.05 and uncorrected p<0.001) among the three different states for both main effect (Figure 7-A) and paired effects (Figure 7-B, C). One prominent cluster was observed in the IC and another in the OR. Post hoc t-tests suggest a change of lag within these two regions between different sensory states, EC vs. EO (Figure 7-B), and EC vs. EOF (Figure 7-C). However, no significant regions were observed when contrasting EO and EOF.

**Figure 7.**
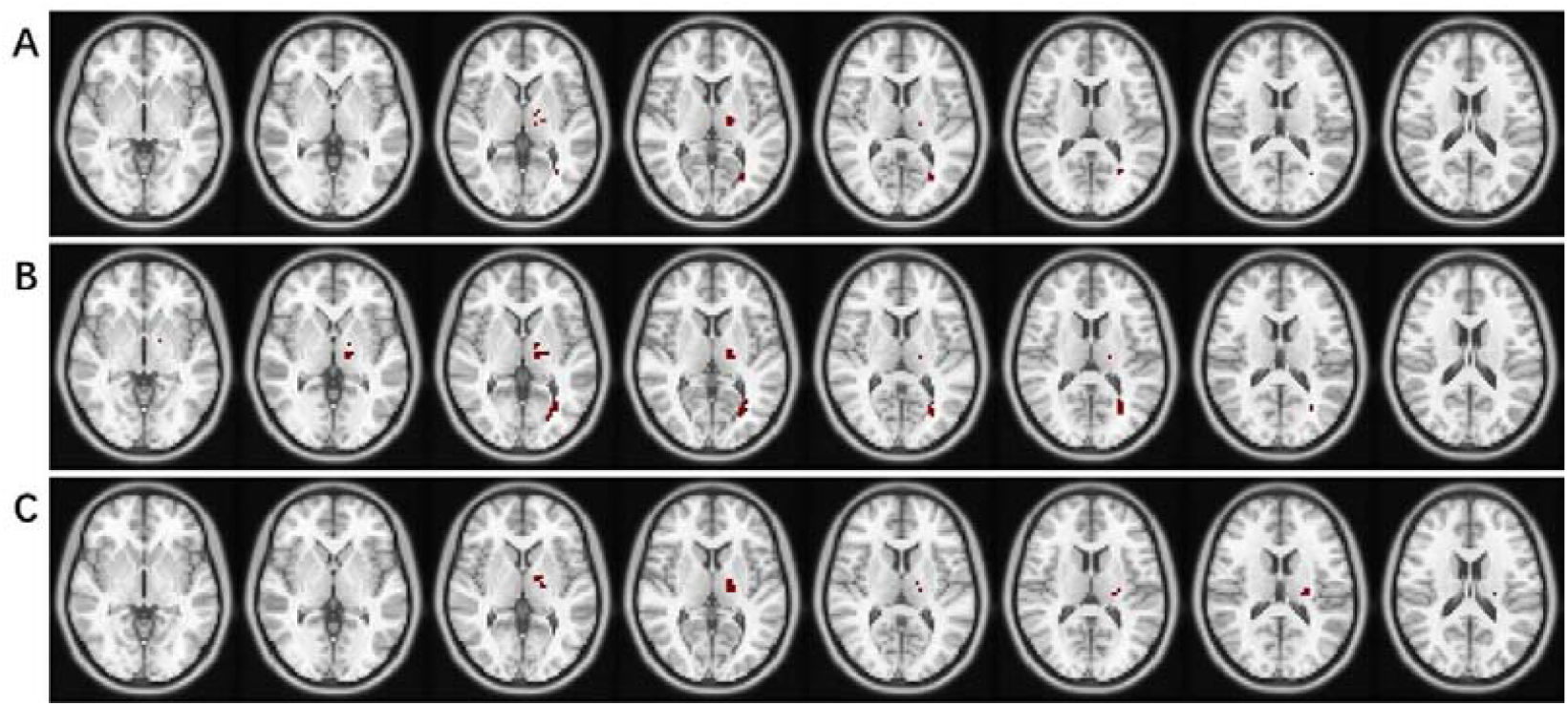
Distributions of white matter voxels with significant effects among different sensory states. (A) Main effect (mixed effect). (B) Paired effect of EC and EO. (C) Paired effect of EC and EOF.

Figure 8 shows the change of latency within regions showing significant effects. As compared with EC, the most prominent change in latency with eyes open was a shift toward later values in the IC, while reverse changes, shifting toward earlier values, were observed in the OR (Figure 6-A, B). Similar changes were also observed by contrasting EC with EOF (Figure 6-A, C). Despite the gross spatial similarity of latency projections seen in Figure 6, the lag values within regions of significant effects changed reliably across different paired sensory states. Specifically, lag values within regions of significant effect changed from -0.46s∼0.44s for EC to -0.16s∼0.17s for EO, and lag values within regions of significant effect changed from -0.46s∼0s (in EC) to -0.14s∼0.07s (in EOF).

**Figure 8.**
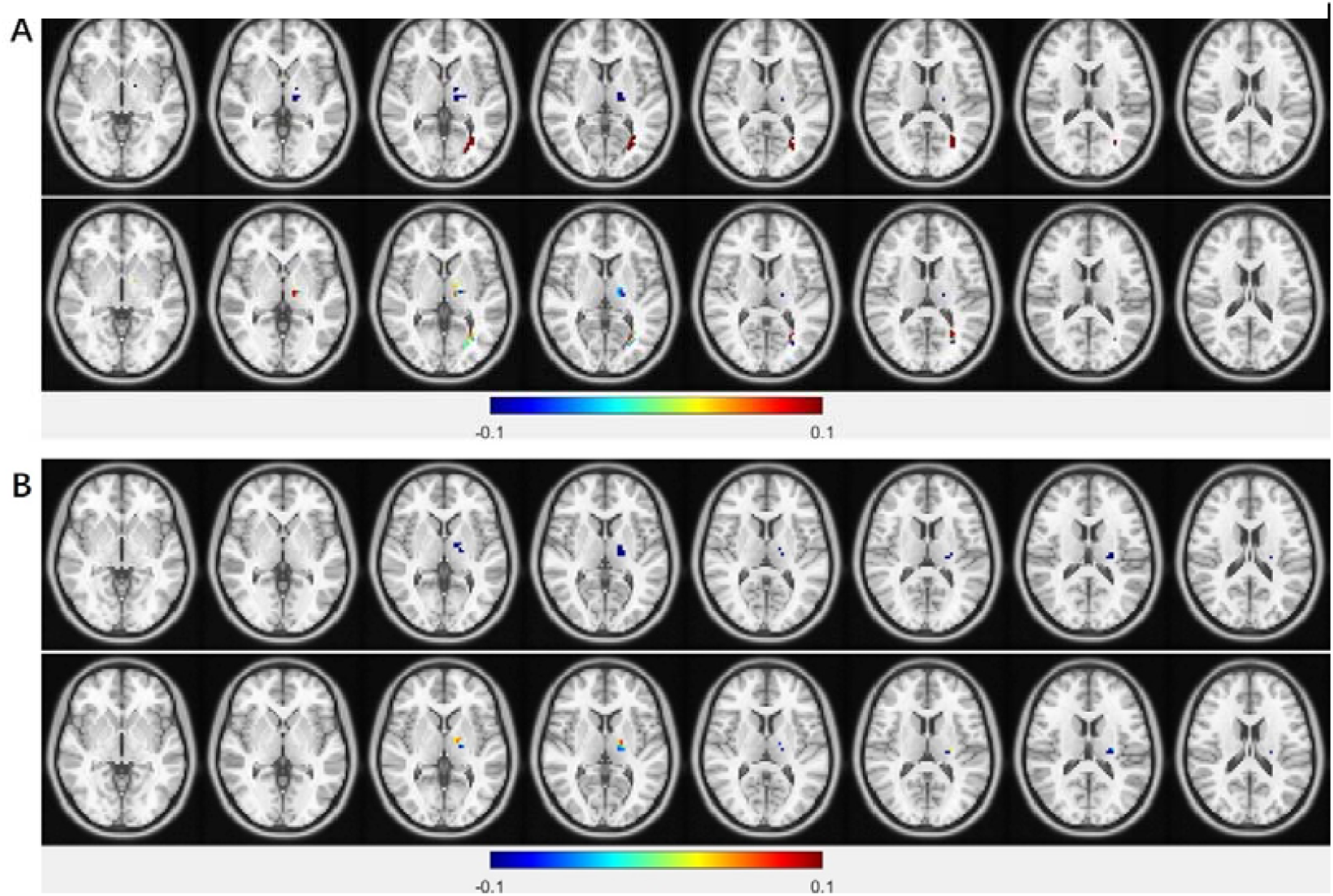
Distributions of latencies in white matter with significant paired effect. (A) EC (top panel) vs. EO (bottom panel). (B) EC (top panel) vs. EOF (bottom panel).

## 4. Discussion

We have described how latency projections [38] derived from time delay matrices were used to investigate the BOLD lag structure within white matter. The consistency of latency projections across ten subgroups consisting of large cohorts of younger healthy subjects attest to the high stability and reproducibility of the lag structure within white matter. Similar stability and reproducibility of lag structure were also observed in gray matter [5]. We also investigated white matter functional networks by applying a clustering technique on group-level functional connectivity matrices. Reproducible functional networks across the ten subgroups were obtained. We further explored how these spatially segregated functional networks are integrated with each other by analyzing inter- and intra-network lead/lag relationships. While some of the networks exhibited reciprocal in-between communication patterns, network-wise lead/lag relations were also revealed. A unidirectional propagation path of BOLD signals was identified from lead/lag analysis of higher resolution functional networks derived from finer grained clustering. We also examined the effects of different sensory states (EO, EC, EOF) to find out whether the latency structure was associated with specific neural inputs. Statistical analysis revealed appropriately localized differences between EO/EC, and between EOF/EC, suggesting that the latency structure in white matter is affected by task-related neural activity.

### 4.1 White matter functional networks

A previous study [21] demonstrated the high reproducibility of bilaterally symmetrical WMFNs. We validated this finding with a much larger dataset. Bilaterally symmetrical WMFNs were also identified in our experiments. However, in the work of Peer et al., the identified WMFNs were organized into three layers (superficial, middle, and deep) with each directly overlaying the preceding one. This spatial overlapping was not observed in this study as our subject-level and subsequent group-level WM masks were generated by highly conservative thresholding (>0.95). As there are lingering concerns that WM BOLD signals may arise from the effects of partial volume averaging with adjacent GM and vasculature, such a tight threshold minimized this potential confound.

Historically, rsfMRI has been extensively used to identify functional networks in the cortex [27][39]. Here we complement previous studies by examining apparent functional networks in white matter. Our results expand beyond previous studies on WMFNs by using a much larger dataset and by quantifying the specific timings of BOLD signals throughout the brain.

### 4.2 Latency of intra- and inter-networks

Intra- and inter-network lead/lag measurements have been analyzed to describe the temporal relationships between spatially segregated resting state networks (RSNs) of gray matter [5]. Studies in human [5][6][40] and mice [41] have reported that BOLD signals propagate through the cerebral cortex along stereotypical spatiotemporal sequences to give rise to the network-level organization. Quasi-periodic patterns (QPPs) of low-frequency activity in BOLD signals were also found to contribute to functional connectivity [42].

In WM, three observations can be derived from network-level latency analyses.

First, some WMFNs exhibited similar intra- and inter-network latency patterns to gray matter (indicated by yellow boxes in Figure 3, 4, 5), suggesting similar spatiotemporal organizational principles governing these networks. For diagonal blocks, this pattern illustrates that each network has early, middle, and late components. With respect to off-diagonal blocks, this pattern represents reciprocal communication between paired networks with no network leading or following the other. Building on a previous study of WMFNs [21], our findings reveal the spatiotemporal organization across RFNs. Future studies could explore factors that affect these results including the influences of different diseases on the functional integration of WMFNs.

Second, temporal orders within some networks (indicated by red boxes in Figure 3) are dominated by the temporal order derived from the latency projection. Each latency block is reordered according to the temporal order obtained from the latency projection. Ideally, if the temporal order within each latency block is consistent with that deduced from the latency projection, which encompasses the global temporal order within white matter, large lags will be uniformly distributed in the top and right position of each latency block. However, these networks turned out to exhibit a reverse temporal order. These regions consist mainly of IC and the anterior part of corpus callosum (BT in Figure 3-D). A preliminary discussion about this was included in Section 4.4.

Third, a dominant propagation direction of BOLD signals within WM can be identified, which suggests that most of the WM BOLD signals propagate along this path. The brain is deemed to be not merely a reflexive but rather a mainly intrinsic model actively interacting with the environment [43]. Functional organizations of the brain are sustained by its intrinsic activities, the spatial coherence of which has been found to transcend different levels of consciousness [44]. From this perspective, the operational model of the brain is to collect and interpret information from the environment, and respond to it whenever necessary. In this interacting progress, the WM is responsible for transferring neural information between the peripheral nervous system and central nervous system (CNS), or within the CNS. The signal propagation along the inferior-superior direction we observed may suggest that during a resting-state, the dominant mode of brain operations is primarily to collect and interpret information from the environment rather than respond to it. Note that the propagation path observed in this work only started from the PLIC instead of lower anatomical regions, e.g. MCP (with pons and medulla included). This may be attributed to the complex and thus plausible disorganized temporal organizations within the MCP. In particular, with high clustering resolution (K=14 in Figure S7), the cerebellum WM was found to be separated from the MCP with the remaining part of the MCP taking the lead in time, which resulted in a unidirectional path pointing from the pons and medulla posteriorly to the cerebellum WM. At this resolution level, the pons and medulla exhibited weakly organized intra-RSN temporal organizations, and also the PLIC seemed to lag the pons and medulla though not overwhelmingly. Further investigations are warranted for clearer understanding of the temporal organizations in the MCP.

It should also be noted that our findings also showed similar observations to previous Electroencephalography (EEG) findings, which has been widely used to measure functional signals and further analyze their propagations in the human brain. For instance, two directions of signal propagations are found in brainstem reticular formation, including descending reticulo-spinal projections for facilitating movement and postural muscle tone with multiple behaviors during wakefulness [45][46] and ascending projections into the forebrain for stimulating cortical activation [47][48][49]. Therefore, in a resting brain, ascending projections should dominate signal propagations presumably because little movement or postural muscle tone take place, which coincides with our findings that the dominant path in BOLD signal propagation is from the inferior to the superior. But further studies are warranted to investigate the directionality and relevance of the dominant path of signal propagations within WM in relation to different brain states, e.g. sleep vs. wakefulness.

Also, we want to point out that network-wise lead/lag relationship was not without precedent in GM [50], though not reported in [5]. Recent studies [51][52][53] converged to demonstrating primary propagation pattern moving from sensorimotor to association cortex, discrepancy between which and [5] may be attributed to the global signal regression (GSR). Future works could also explore the influence of GSR on the derived spatiotemporal organizations.

### 4.3. Influences of different sensory states on white matter latency structure

Given the observed effects of different sensory states on latency, it is reasonable to infer that these effects are potentially of neurophysiological origin. Though latency projections from different sensory states exhibited similar tendency of increasing lag values along OR, lag values changed significantly when contrasting EO with EC, and also EOF with EC. While comparisons between EO and EOF showed no significant changes in OR, WM BOLD signals in OR show close associations with visual tasks [54][17][13]. Ding et al. also found that visual stimulations significantly enhance the correlation between gray matter and OR [17], which may explain the lower estimated lag values in OR. The change of lag values due to different sensory states may suggest a different extent of engagement of OR, in which a smaller lag value in OR corresponds to more active engagement of OR in modulating visual information. Likewise, explanations may extend to the IC as well, as it consists of both ascending and descending fibers. This structure, situated close to the thalamus, routes information up to the corona radiata and down to the pons. If the IC is more actively engaged in a task, the time delays within the affected region may be shortened.

### 4.4. Latency structure of white matter in relation to vascular physiology

BOLD signals result from variations of regional blood concentrations of oxy- and deoxy-hemoglobin. Hence the latency structure may be determined by the anatomy of the venous drainage system within white matter. Previously in gray matter, the venous drainage system was shown to be not relevant to the latency structure [5]. Indeed, for gray matter, the superficial venous system drains into the superior sagittal sinus from an inferior to superior direction, while the superior sagittal sinus drains the venous blood from superior to inferior regions. However, the distribution of the lag values does not follow the flow direction of the superficial venous drainage system, which therefore rules out the possibility that the latency structure merely reflects the vascular architecture. Likewise, generally in white matter, the deep venous drainage system removes venous blood from anterior (via the inferior sagittal sinus, the internal cerebral vein, and the basal vein) to posterior (the straight sinus, the great cerebral vein) directions. Again, the distribution of the lag values within white matter are inconsistent with this flow. From our analyses a unidirectional path of BOLD signal propagation was revealed, from inferior to superior. Moreover, the lag values in the IC and the OR were significantly associated with different sensory states, which is not explicable in terms of the deep venous drainage system.

As we based our study on fMRI BOLD signals, we cannot exclude hemodynamic effects on latency projections. It was revealed by a previous study that bundle-level hemodynamic delays between superficial and deep WM could be of seconds, wherein deeper regions in WM showed larger lags [13]. A reasonable extension of this finding is that hemodynamic response functions in WM shared different delays even among neighboring voxels. This intrinsic property may give rise to the observed latency projection. Actually, our findings of latency structure exhibited similar tendency of delays especially in the distributions of lead/lag values, wherein large lags (positive values) were mostly located in deep WM including four corners near the ventricle, OR, and CR. This lead/lag distribution may be attributed to the arterial structure for blood supply of WM. Despite that BOLD effect is directly related to the concentrations of deoxyhemoglobin in venous blood [55], over-supplied oxygenated blood due to uncoupled regional cerebral blood flow (CBF) and cerebral metabolic rate of oxygen consumption [56][57] was provided via arteries. The arterial supply of WM consists of subcortical arteries [58], for blood supply of subcortical WM, and medullary arteries [59][60], for blood supply of deep WM. Subcortical arteries penetrate straight through the cortex to supply for subcortical WM, while medullary arteries also penetrate straight through the cortex but to supply for deep WM with longer time of delay than that of subcortical arteries. This arterial structure may be the physiological basis of the observed latency structure. On the other hand, the regional variations of lags in IC and OR may be attributed to the changes of regional neurovascular coupling due to different underlying neural activities across different sensory states, leading to early shifting of localized BOLD signals from eye closed to eye open. Since both arterial supplies of GM and WM branch off from pial arteries, it lingers that BOLD effect from WM may be the concomitant effect of GM, regionally neuronal activations of which will induce retrograde propagation of vasodilation to the area of pial arteries [61], thus yielding the BOLD effect of WM in return. However, it should be noted that the increases in CBF that are evoked by neural activity are highly site-specific [62]. For example, in the olfactory bulb, stimulated specific glomeruli produce increased flow in capillary involving the activated glomerulus, but not capillaries that are even less than 100 µm away in quiescent areas [63]. Hence, neural activity from GM cannot induce concomitant BOLD effect in WM. In fact, it has been shown that changes in brain state from eye closed to eye open were reflected in different extents of increased FDG uptakes in GM and WM [64], suggesting actual changes in neural activity between these two brain states. To provide further evidence, a recent study [65] based on simultaneous MRI and PET of healthy human subjects revealed a correlation between functional and metabolic measures in WM, suggesting that functional involvement of WM in cortical processing was associated with metabolic demands of underlying neural activities. Compared with that in GM, WM has been found to contain a significantly higher glia-to-neuron ratio [66]. Also, a recent study argued that oligodendrocyte activities make up a significant portion of the metabolic demands in maintaining membrane potentials for proper functioning [67] or supporting metabolic demand of neurons [68], which may be the primary source of metabolic demands. Therefore, the latency structure may be related to the structure of arterial supply of WM and regional variations of lags across brain states may be attributed to different underlying neural activities.

However, we also need to note that the arterial structure alone, if attributable, could not explain the total variations of the latency matrix. In fact, if we applied principal component analysis (PCA) on the latency matrices of the ten subgroups from GSP dataset (refer to Figure S8 in supplementary material for PCA results), principal components with the largest two eigenvalues, explaining in average ∼45% and ∼20% variances of the latency matrix, respectively, were identified. This suggested that there existed more than one latency processes contributing to the latency matrices reflecting voxel-wise temporal organizations within WM.

The networks showing reversed inter-network against intra-network temporal ordering (red boxes in Figure 3, RPIC in Figure 4, and also white matter in cerebellum (network 3) in Figure S7) was consisted mainly of IC (the network RPIC, which also includes other part of IC, is named according to the largest bundle it contains). Here we provide a speculative example of this observation. By performing latency analysis among voxels within RPIC, it can be seen that the latency projection of RPIC are well-ordered (top panels in Figure S9-A). We could further divide the network into two regions as seeds (middle and bottom panels in Figure S9-A), an early seed (defined by thresholding the above latency projection at lags < 0) and a late seed (defined by thresholding the above latency projection at lags > 0). Seed-based latency maps were computed to evaluate the relative lags between each seed and the remaining voxels within WM (top and middle panels in Figure S9-B). It can be seen from the difference map (bottom panels of Figure S9-B, paired-sample ttest, p < 0.05 with FDR correction) that early region of RPIC showed smaller lag with respect to the most remaining voxels of WM than late region, which suggested a propagation pattern when early region initiated not only the intra-network propagation to late region within RPIC but also the inter-network propagation to other networks (top panel in Figure S9-C wherein E and L represent early and late regions of RPIC, respectively, and T1∼T3 represents three nodes from superior networks). A latency matrix of the propagation pattern could be created (middle panel in Figure S9-C; the lead/lag values among T1∼T3 are left blank since these values did not affect the derived relative timing between E and L), which is a simplified version of the observed results in Figure S9-A,B (Note the larger lags of the node T1∼T3 with respect to node L than to node E). From this latency matrix, a correct relative timing between E and L (green timing line in bottom panel in Figure S9-C) can be derived if we used only the lead/lag relationship in green rectangle, i.e., to include their temporal orders from within RPIC. If the temporal orders between nodes E and L are derived in a global sense (the rectangle in the middle panel of Figure S9-C), a reversed temporal order against the intrinsic one is derived (red timing axis in bottom panel in Figure S9-C). Speculatively, this observation may be related with multisensory integration [69][70] concerning the transmission and integration of different sensory signals, e.g. information from E and L is required in T1∼T3, but L should be activated by E before the propagation from L to T1∼T3 can be started (Figure S9-C). In fact, IC may possess the specialty to show this type of temporal organization since both ascending and descending axons densely abound in IC to connect the cerebral hemispheres with subcortical structures, the brainstem, and the spinal cord. While the functional networks derived from K-means failed to distinguish these two types of axons, a tractography based parcellation method may help to identify the ascending and descending targets, thus promoting a better understanding of this type of temporal organization in a physiological sense.

### 4.5. Acquisition length in relation to reliability of latency projection

Reliable group-level RSFC can be obtained with acquisition length of 5-13 minutes, wherein reliability increased with acquisition length [71]. It should be noted that, however, longer time of acquisition tends to incur greater imaging artefacts, due to e.g. head motion or cognitive instability, which eventually compromised the benefit brought by long acquisition time. Hence, sampling error can be reduced by averaging across several scans rather than a very long duration scan. A detailed investigation of the influences of factors including correlation magnitude, data length, and temporal resolution, on time delay estimation can be found in [34], though using either surrogate time series or multiple-session individual real data [34][72]. In spite of the sampling error in estimated lags, latency analysis based on current method [5] could give rise to differences between groups [73][74][75] using typical scanning parameters (e.g. TR = 2000ms, acquisition length = 8min or 10min). Here we evaluate the influences of acquisition length, along with the sample size, on the reliability of group-level latency projection. We selected the first subgroup from the GSP dataset and performed latency analysis but with various acquisition lengths, ranging from 1-6 minutes, at various sample sizes (n=10, 20, 40, 80). For each combination of the acquisition length and sample size, latency analysis was repeated 10 times. Pearson’s correlations were evaluated between the original latency projection from this group and the latency projection from each of the combinations. As clearly shown in Figure S10, the reliability increases with the acquisition length at all sample sizes; when the acquisition time was beyond 4 minutes, the curves tend to be flat, indicating that additional benefit brought by acquisition time longer than 4 minutes tends to be small.

### 4.6 Clinical Implications of WM Latency Analysis

WM, especially the corpus callosum, has a causal role for building up inter-hemispheric FCs [76] by transducing neuronal information in myelinated axons. The unique and pivotal role of WM has important implications to studies of brain development, aging and many neurodegenerative diseases. For instance, in multiple sclerosis the myelin that covers neuronal fibers is impaired, so signal communications between brain regions are thus less efficient. In such a clinical scenario, analyzing the WM latency may assist in evaluating the altered spatiotemporal organization within WM so that possible breakdowns of intra- and inter-RSN integrations may be identified. Similarly, in Alzheimer’s Disease (AD) which involves degenerations of myelin that are potentially associated with age-related deficits in memory [77], analyzing the WM latency might add to the repertoire of potential biomarkers for early diagnosis of AD.

## 5. Conclusion

In summary, based on a large fMRI dataset acquired from youngsters, we investigated in this study resting state WM latency structure, which represents apparent propagations of fMRI BOLD signals. We found there are lead/lag relationships among WM networks, and different sensory states entail different latency structures. Also, we have identified a unidirectional path of signal propagations within the brain WM, which is suggestive of a special mode of brain functional operations. This study offers a new perspective for looking into the temporal organization of the brain particularly WM, which may have important implications to studies of brain development, aging and neurodegenerative diseases.

## Author Contributions

Bin Guo conceived the project and research approach. Bin Guo designed the methods. Bin Guo implemented the methods and performed the data analysis. Bin Guo processed the data. Bin Guo wrote the paper; Fugen Zhou, Muwei Li, John C. Gore, reviewed and edited it.

## Acknowledgements

This work was supported by the National Institutes of Health(NIH) grant R01 NS093669 (J.C.G) a nd R01 NS113832 (J.C.G). Also, we sincerely thank Dr. Zhaohua Ding for his insights in our discussion.

