## supplementary material for "Latency structure of BOLD signals within white matter in resting-state fMRI"

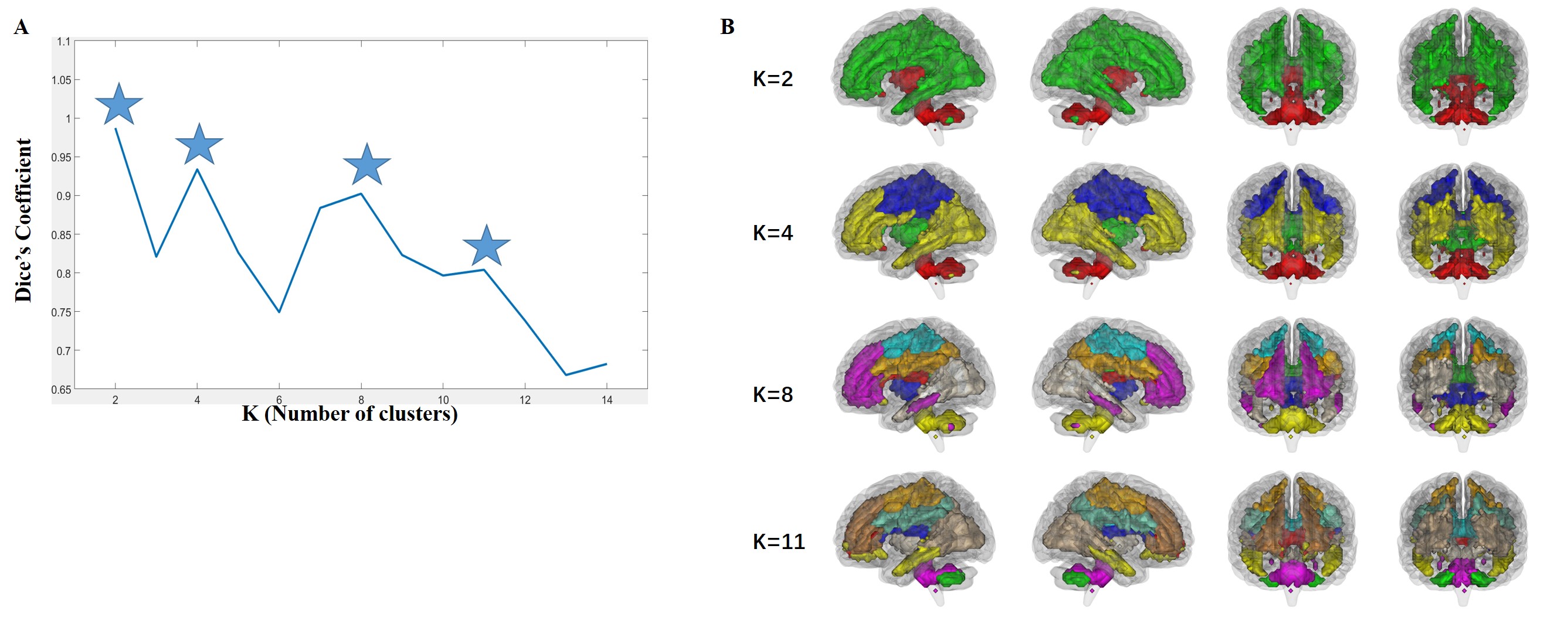

Figure S1 Variations of clusters obtained from white matter correlation matrix with values of K. (A) Average Dice Coefficients over the 10 subgroups, with asterisks denoting local maxima. (B) From top to bottom are 2, 4, 8 and 11 functional networks clustered from white matter correlation matrix. Highly bilaterally symmetrical functional networks are observed.

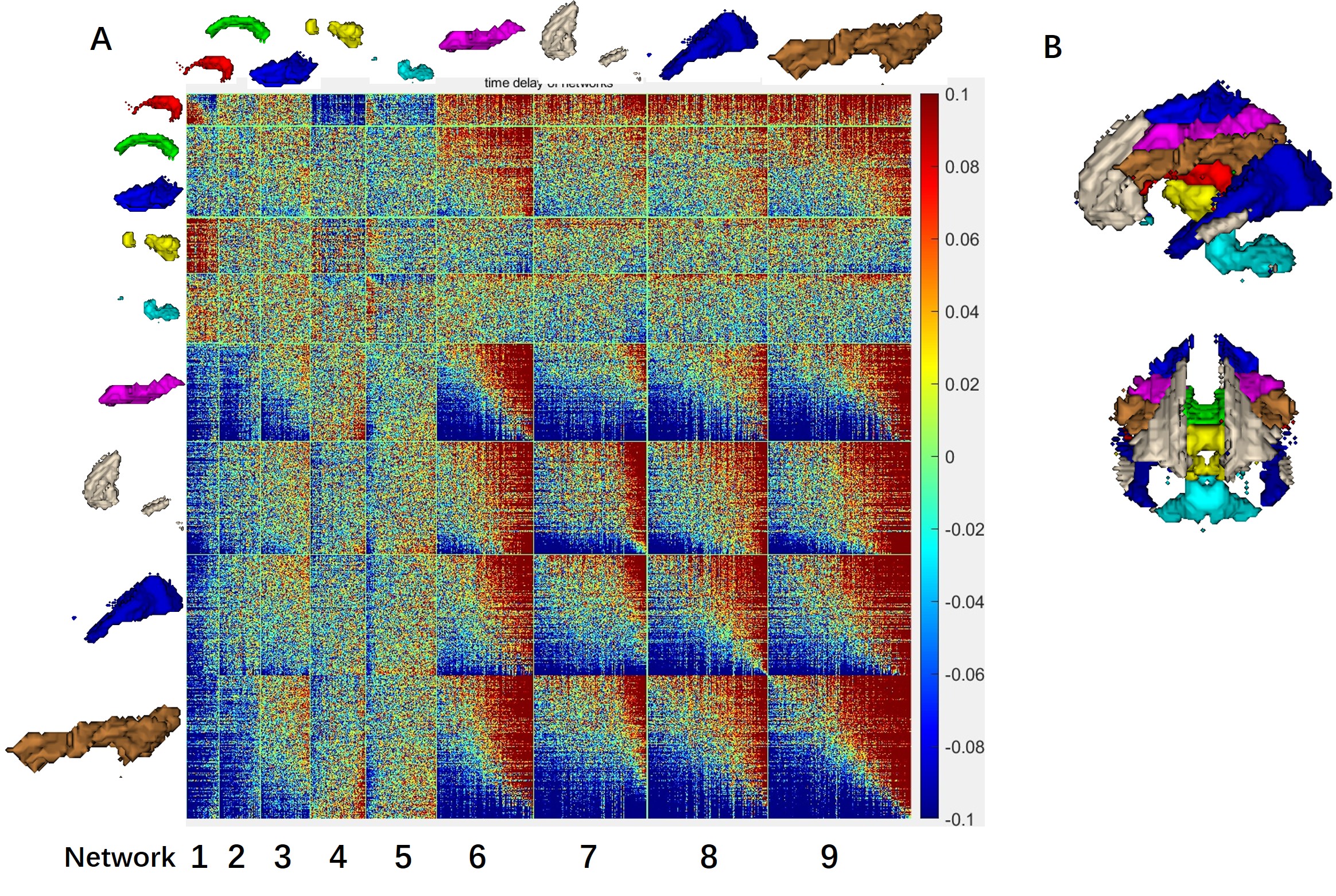

Figure S2 Intra- and inter-network lead/lag analysis on nine clusters. (A) The intra- and inter-network lead/lag array. (B) The left and front view of the clustering results. Blocks referred to in the Section of Discussion are outlined in different colors.

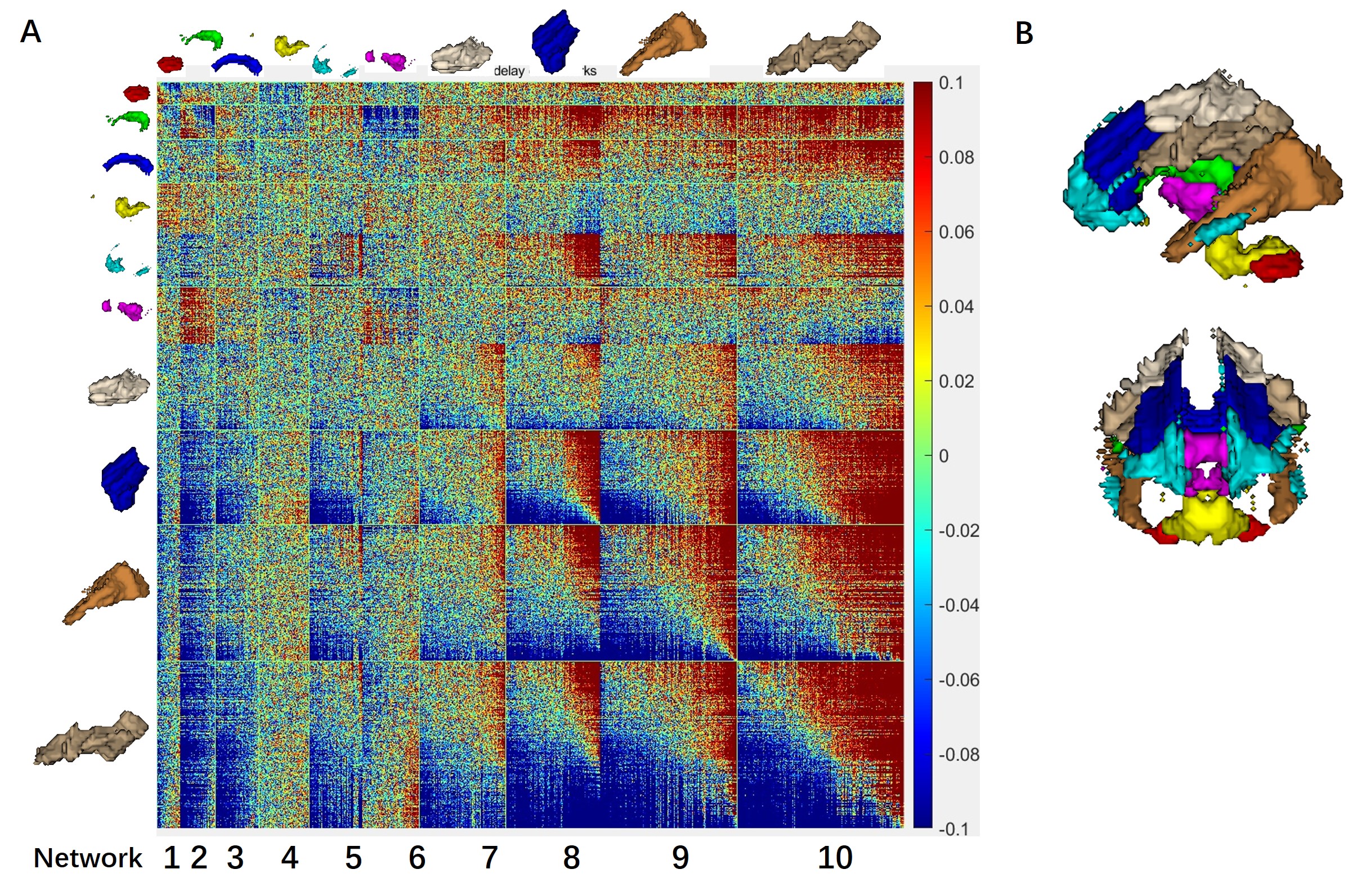

Figure S3 Intra- and inter-network lead/lag analysis on ten clusters. (A) The intra- and inter-network lead/lag array. (B) The left and front view of the clustering results. Blocks referred to in the Section of Discussion are outlined in different colors.

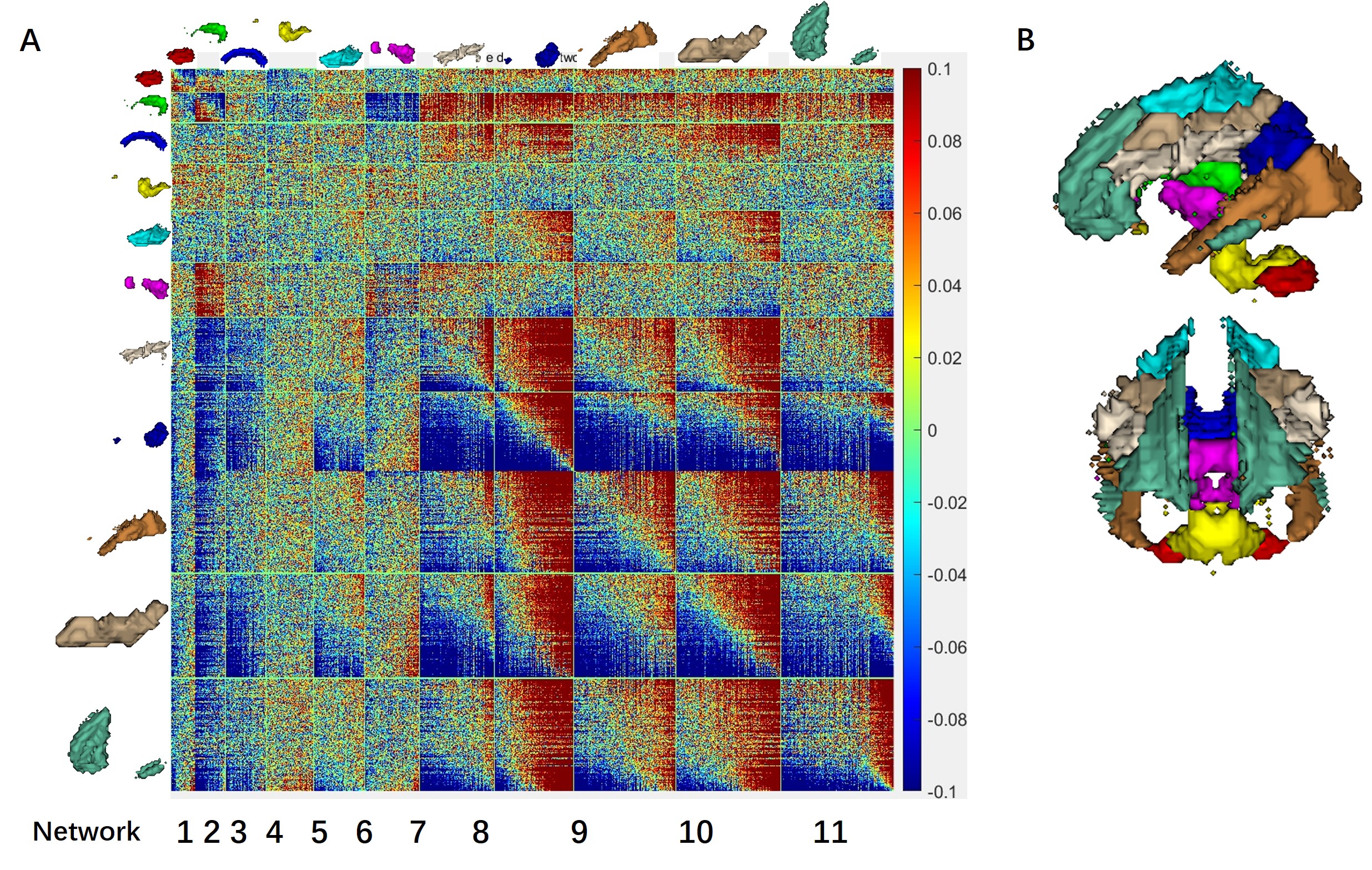

Figure S4 Intra- and inter-network lead/lag analysis on eleven clusters. (A) The intra- and inter-network lead/lag array. (B) The left and front view of the clustering results. Blocks referred to in the Section of Discussion are outlined in different colors.

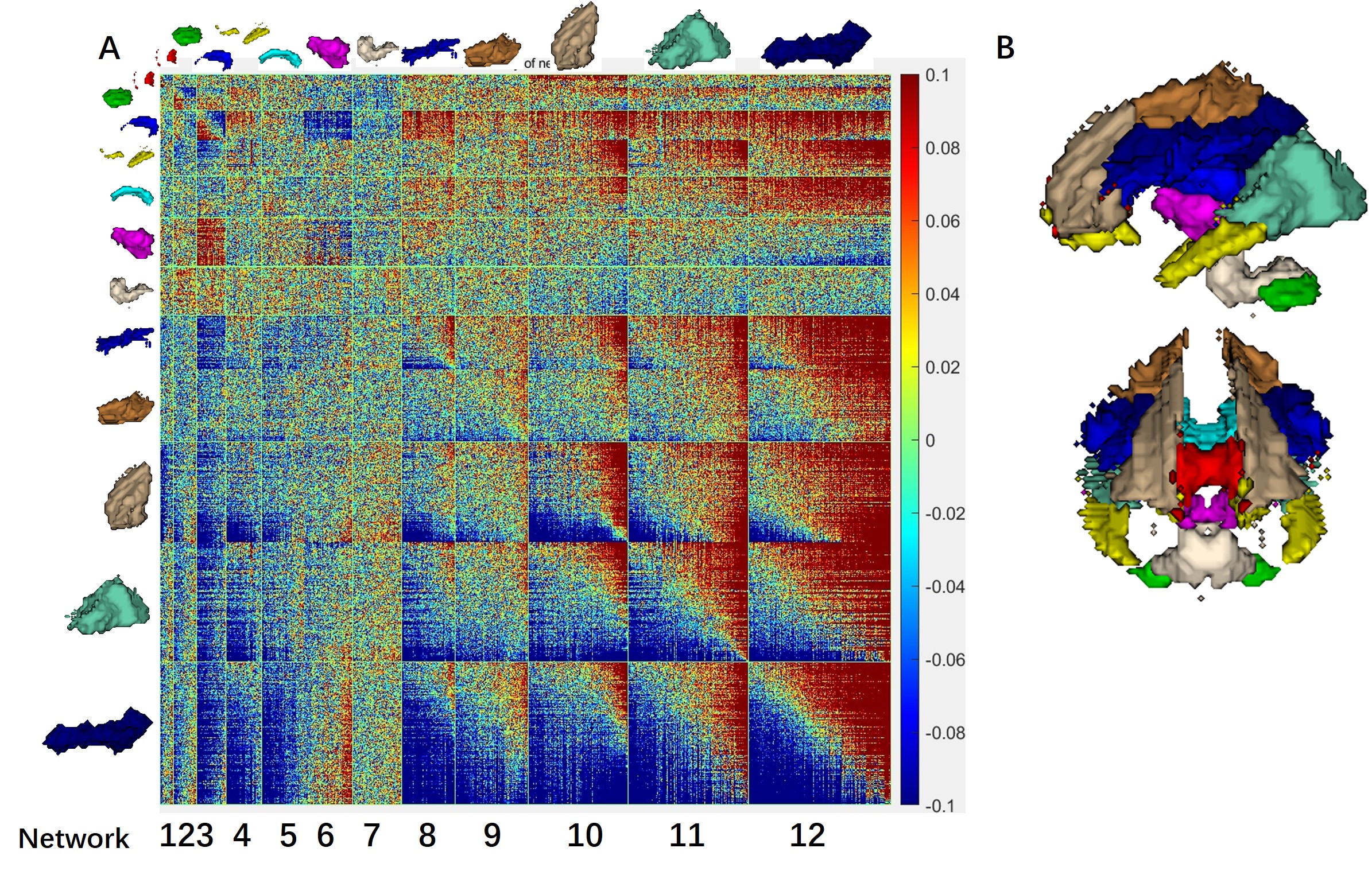

Figure S5 Intra- and inter-network lead/lag analysis on twelve clusters. (A) The intra- and inter-network lead/lag array. (B) The left and front view of the clustering results. Blocks referred to in the Section of Discussion are outlined in different colors.

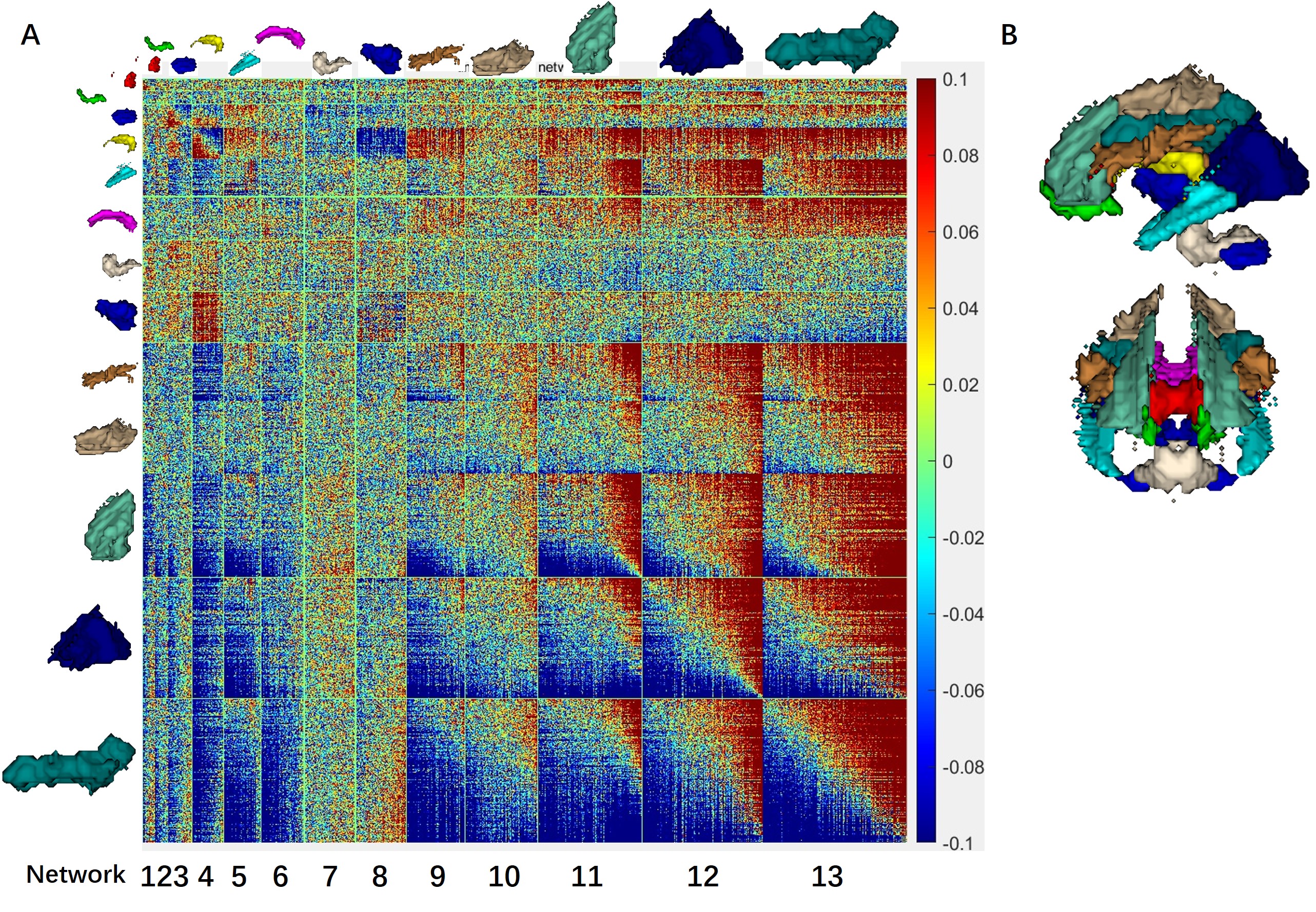

Figure S6 Intra- and inter-network lead/lag analysis on 13 clusters. (A) The intra- and inter-network lead/lag array. (B) The left and front view of the clustering results. Blocks referred to in the Section of Discussion are outlined in different colors.

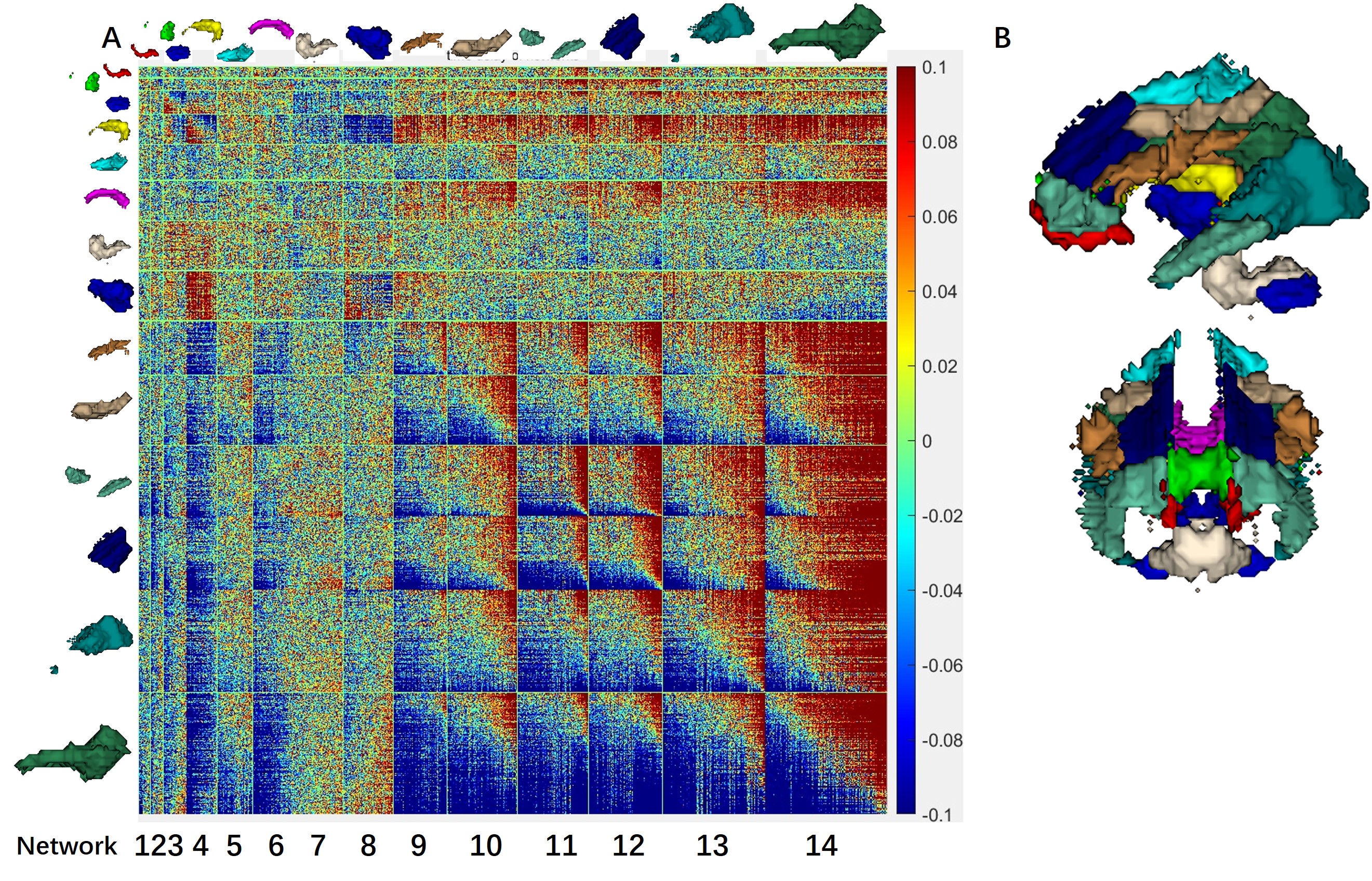

Figure S7 Intra- and inter-network lead/lag analysis on 14 clusters. (A) The intra- and inter-network lead/lag array. (B) The left and front view of the clustering results. Blocks referred to in the Section of Discussion are outlined in different colors.

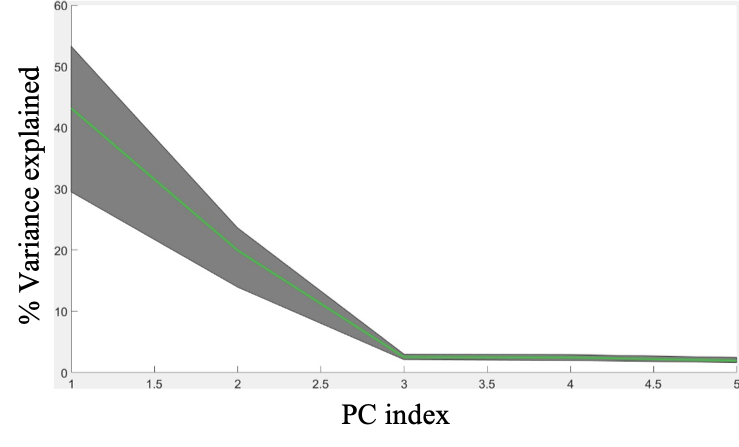

Figure S8 PCA analysis on the latency matrix of each subgroup from GSP dataset. Regions shaded in gray represent the eigenspectrum derived from the ten groups. The green line indicated the average eigenspectrum over all subgroups.

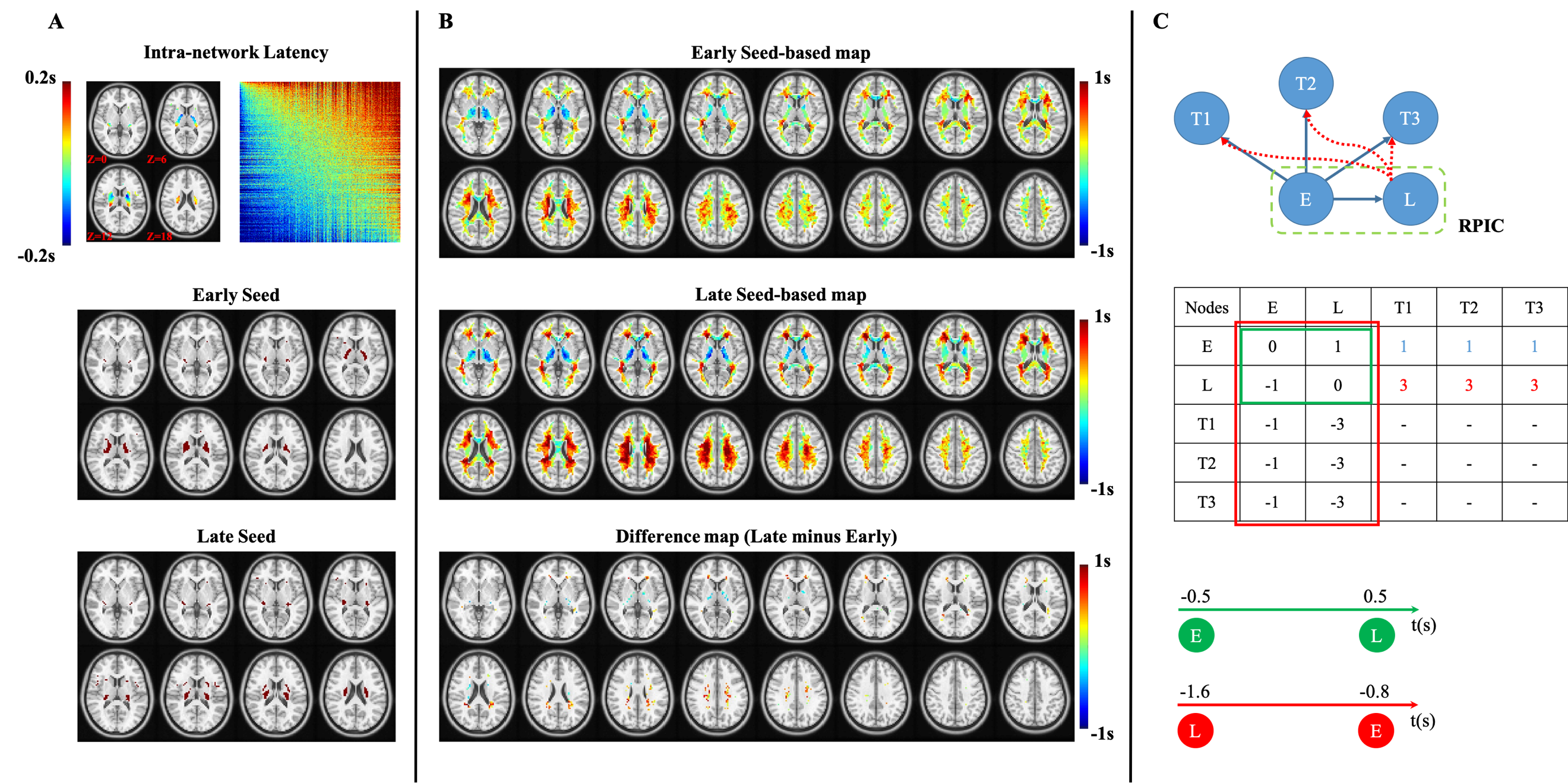

Figure S9 Illustrations of reversed temporal organizations within RPIC between intra- and inter-network derivation. (A) Top-Left: Intra-network latency projection of RPIC, derived using voxels from RPIC only; Top-Right: Intra-network latency matrix derived using voxels from RPIC only wherein well-ordered temporal components were identified. Middle: Regions containing early timing voxels from the latency projection above are regarded as early seed; Bottom: Regions containing late timing voxels from the latency projection above are regarded as late seed; (B) Top: Early seed-based latency map, reflecting the relative timing between early seed and each of the remaining voxels in WM; Middle: Late seed-based latency map, reflecting the relative timing between late seed and each of the remaining voxels in WM; Bottom: Difference map between late seed-based latency map and early seed-based latency map (late minus early). Voxels showing statistical significance (paired-sample ttest, p < 0.05 with FDR correction) are displayed. (C) Top: A simplified illustration of the temporal organizations in (A), (B), wherein E indicates the early seed, L the late seed, and T1~T3 represent three nodes from the superior WM. Node E initiates the propagations from not only node E to node L but also to T1~T3 (indicated by blue arrows). Propagations from node L to T1~T3 require larger lags (indicated by red dashed arrows) than node E; Middle: Latency matrix of the nodes in the top panel. Green rectangle indicates regions used for derivation of intra-network latency projection while red rectangle inter-network latency projection. Bottom: Relative timing between node E and node L when the temporal orders are derived from intra-network manner (time axis in green) or inter-network manner (time axis in red).

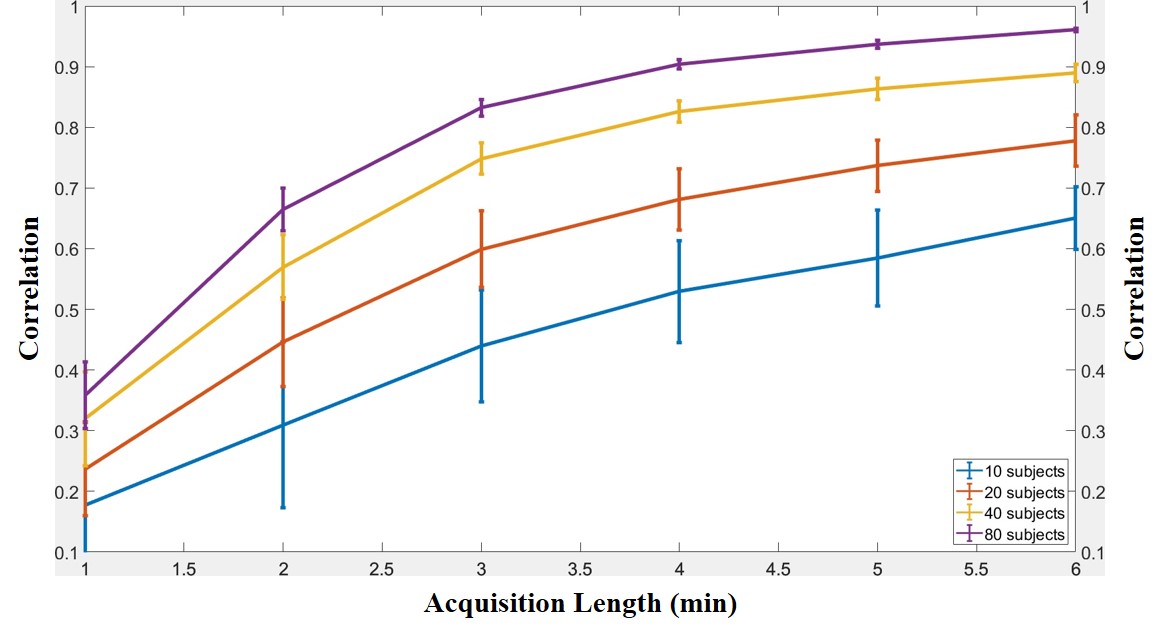

Figure S10 Correlations between the original latency projection of the first subgroup and each of the latency projection derived from different acquisition length, starting from 1 minute to 5 minutes, of the same subgroup. Different curves indicated the results from different number of subjects.
